# Conbase: a software for unsupervised discovery of clonal somatic mutations in single cells through read phasing

**DOI:** 10.1101/259994

**Authors:** Joanna Hård, Ezeddin Al Håkim, Marie Kindblom, Åsa K. Björklund, Ilke Demirci, Marta Paterlini, Pedro Reu, Bengt Sennblad, Erik Borgström, Patrik L. Ståhl, Jakob Michaelsson, Jeff E. Mold, Jonas Frisén

## Abstract

Here we report the development of Conbase, a software application for the identification of somatic mutations in single cell DNA sequencing data with high rates of allelic dropout and at low read depth. Conbase leverages data from multiple samples in a dataset and utilizes read phasing to call somatic single nucleotide variants and to accurately predict genotypes in whole genome amplified single cells in somatic variant loci. We demonstrate the accuracy of Conbase on simulated datasets, *in vitro* expanded fibroblasts and clonally *in vivo* expanded lymphocyte populations isolated directly from a healthy human donor.

## Background

Single cell DNA sequencing has substantial untapped potential for understanding genomic diversity in both healthy and disease states and for reconstructing phylogenetic relationships of individual cells^1,2^. However, current methods for single cell DNA sequencing typically require a whole genome amplification (WGA) step to yield enough DNA for sequencing, which is widely recognized to introduce substantial technical artifacts including allelic dropout, amplification bias, as well as amplification errors leading to false negative and false positive variant calls^3,4,5,6^. Additionally, in traditional whole genome sequencing data obtained from bulk genomic DNA, the error rate of raw variant calls is 1 in 10-15kb, predominantly resulting from alignment artifacts caused by failed realignment and an incomplete reference genome with respect to the genome of the donor^7^. These are likely to be major concerns for single cell variant calling. Given that somatic mutations in normal (non-malignant) cells are generally estimated to be infrequent (approximately three somatic mutations per cell division^2^), the expected number of false positive variants far exceeds the predicted number of true somatic mutations.

In WGA data, expected observations include sites displaying reads originating from both the maternal and the paternal alleles, as well as sites displaying reads originating from only one of the two alleles (due to allelic dropout)^3,4,5,6^. Sites displaying reads originating from only one allele may result in falsely predicted reference genotypes (false negative genotype calls) if dropout occurs only for the mutated allele. In addition, sites may be covered by reads derived from multiple locations in the genome (for instance due to failed realignment and structural variation that differ between the reference genome and the genome of the donor), or from the same location in the genome, where a subset of reads contain a mismatch against the reference genome (due to WGA errors)^3,4,5,6,7^. In the absence of additional information, such observations can be indistinguishable from true somatic mutations and will result in falsely predicted alternative genotypes with support for a non-reference base (false positive genotype calls). Taken together, alignment artifacts, amplification errors and allelic dropout result in false positive and false negative genotype calls, hampering the use of variant calling to define phylogenetic relationships at the single cell level^4,5^.

In order to circumvent these issues, we developed a computational strategy for the unsupervised discovery of somatic single nucleotide variant sites (sSNVs) in single cell whole genome sequencing data, including accurate genotyping of the individual single cells independently of the global rate of allelic dropout. Conbase is a multistep algorithm that <u>con</u>firms the allelic origin of <u>base</u>s through read phasing, by using the abundant signal from germline single nucleotide variants (gSNVs) across the genome. The discovery of sSNV sites is based on analysis of observed haplotype concordance within individual single cells, across the dataset and in an unamplified bulk sample. By further exploiting the phasing information, locus specific allelic dropout is determined per sample individually, enabling exclusion of false negative genotypes resulting from dropout of the mutated allele.

We demonstrate the specificity of Conbase on simulated data and two different real datasets containing single cell DNA libraries from healthy human cells prepared using different WGA techniques. Both datasets exhibit high rates of allelic dropout, and one exhibits in particularly high error rate. One dataset was generated from CD8^+^ T cells, making it possible to evaluate the specificity of variant calling in real data isolated from healthy human subjects, since the true clonal relationships can be confirmed by parallel analysis of rearranged T cell receptor genes. Using this dataset, we perform extensive evaluation of variant calling output and comparative analysis with the recently described single cell variant caller Monovar^4^. Indeed, Conbase outperforms Monovar in calling true sSNVs in real world data obtained from WGA amplified single cells isolated from a healthy human donor.

In summary, we demonstrate the effectiveness of Conbase for identifying true sSNVs in single cell DNA sequencing libraries of varying quality from biologically relevant populations of human cells. We believe that Conbase will be an increasingly valuable tool for applications ranging from phylogenetic analysis of single eukaryotic cells on the basis of acquired sSNVs to the characterization of the mutational landscape of single cells in healthy and diseased tissues.

## Results

Overview of Conbase variant calling

Allelic dropout and errors caused by amplification errors and alignment artifacts can result in false negative and false positive genotype calls, respectively (illustrated in Figure 1). Conbase circumvents these problems by integrating phasing and analysis of observed haplotype concordance during variant calling (Figure 1). Phasing putative sSNVs to gSNVs allows for the determination of maternal or paternal origin of variants, because true sSNVs are expected to be observed only on either the maternal or the paternal allele in the population of cells (Figure 1). The allele that harbors a variant in mutated samples is here defined as the informative allele. The informative allele is distinguished from the non-informative allele by the base observation in gSNVs present in the same sequenced molecule (Figure 1). The genotypes of samples that only display reads originating from the non-informative allele are unknown (Figure 1). False negative variant calls resulting from allelic dropout, are eliminated by requiring for a sample to have reads originating from the informative allele in order to be assigned a genotype (Figure 1). False positive variant calls result from amplification errors, as well as systematic errors including alignment artifacts, structural variation and errors in the reference genome. These are to a large extent excluded by analyzing multiple observed data during variant calling and genotyping, including monitoring observed haplotypes in putative sSNV loci (Figure 1) as well as maximal expected read depth in an unamplified bulk sample and the density of mismatches against the reference genome in the region in which variant calls are present (Methods section: Algorithm description, gSNV filtering).

**Figure 1.**
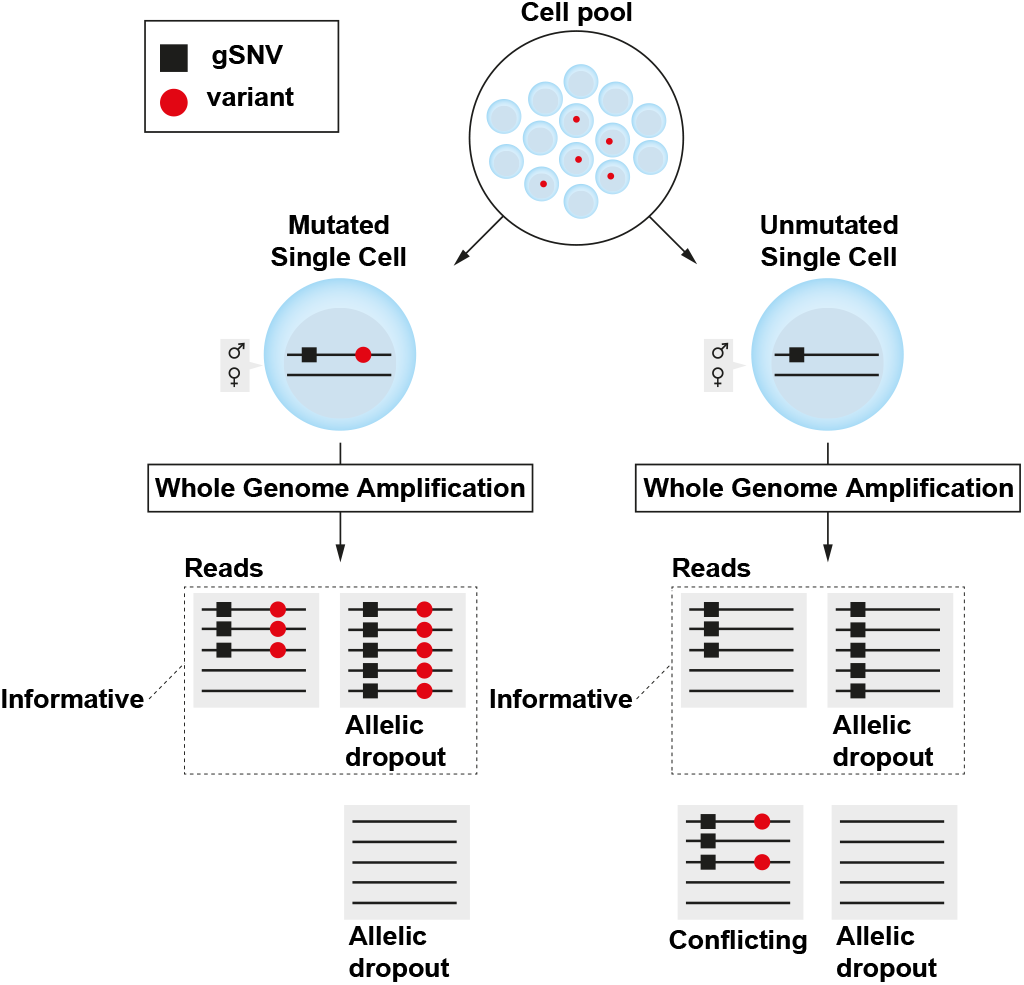
Somatic mutations are present in a subset of a population of cells and can be identified by DNA sequencing of whole genome amplified single cells. Whole genome amplification may result in allelic dropout, which in turn may result in false negative variant calls if dropout has occurred of the mutated allele. False positive variant calls may arise from amplification errors or alignment artifacts by read molecules with high sequence similarity, resulting in conflicting haplotype observations. Conbase circumvents these problems by determining locus specific allelic dropout individually per sample and analyzes concordance of the observed haplotypes across the cell population.

### Performance evaluation on simulated data

To evaluate the specificity of Conbase, we generated synthetic mapped read file datasets of samples with known genotypes to study how allelic dropout and errors affect the ability of Conbase to correctly predict genotypes (Figure 2, Methods). We chose to evaluate the performance of Conbase in comparison with the recently described variant caller Monovar, which was developed specifically to identify SNVs in single cell libraries^5^. Monovar models allelic dropout rate and WGA error rate to calculate likelihood scores for the predicted genotypes, and demonstrates superior performance when compared to conventional tools for variant calling designed to analyze bulk genomic sequencing data^4^. To study the ability of Conbase to predict correct genotypes in the presence of allelic dropout, we synthesized samples with increasing rates of allelic dropout, by random removal of all reads originating from one of the two alleles defined by either the reference base or the alternative base in a neighboring gSNV present in the same molecule. In all samples affected by allelic dropout, the true genotype was the alternative genotype in all sites, thus displaying support for a nonreference base in the simulated variant site before allelic dropout was applied (Figure 2a). In sites where the true genotype is the alternative genotype, allelic dropout may however result in false negative genotype calls if dropout of the mutated allele has occurred (Figure 2a). As such, we could determine the fraction of false negative genotype calls by the number of sites in which the reference genotype was predicted. In samples affected by allelic dropout, no sites with false negative genotypes (reference genotype) were present in the Conbase output (Figure 2b, Supplementary Figure 1). In the output from Monovar, we observed sites where the reference genotype was incorrectly predicted (Figure 2b, Supplementary Figure 1). The fraction of false negative genotype calls per sample in the Monovar output correlates with the rate of allelic dropout (Figure 2b). At a 100% allelic dropout rate, there was an equal probability of Monovar to assign the reference genotype or the alternative genotype to sites where the true genotype was the alternative genotype (Figure 2b). Sites that were incorrectly assigned the reference genotype by Monovar, were classified as ‘missing data’ by Conbase (Figure 2a,c, Supplementary Figure 1).

**Figure 2.**
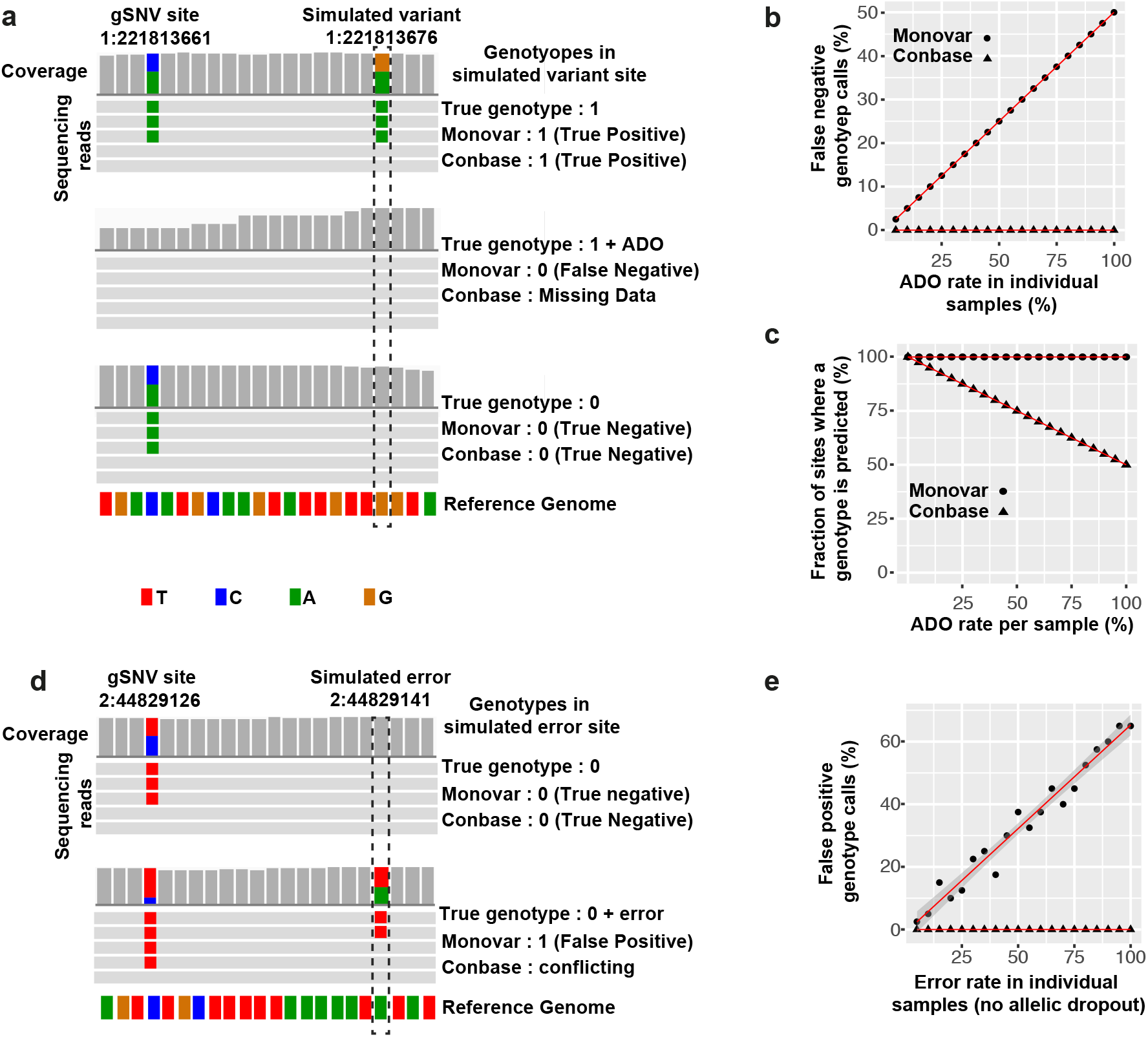
(a) Simulated sequencing read data visualized in Integrative Genome Viewer. Top panel represents a mutated sample with reads from both alleles. Middle panel represents a mutated sample affected by allelic dropout of the mutated allele. Bottom panel represents an umutated sample with reads from both alleles. (b) False negative genotype calls in variant calling output from Conbase and Monovar in a dataset where an increasing fraction of sites per sample is affected by simulated allelic dropout. Each data point represents one sample. (c) The fraction of sites in which a genotype was predicted by Conbase and Monovar in the presence of increasing levels of allelic dropout (d) Simulated sequencing read data visualized in Integrative Genome Viewer. Top panel represents an unmutated sample with reads from both alleles. Bottom panel represents an unmutated sample affected by errors, resulting in support for conflicting genotypes. Sites where conflicting genotypes are observed, are by default filtered out as low confidence variant sites by Conbase. (e) False positive genotype calls in variant calling output from Conbase and Monovar in samples where an increasing fraction of sites per sample is affected by errors.

Amplification errors and alignment artifacts may result in false positive variant calls (Figure 2d). In the population of samples, systematic errors caused by for instance incorrectly mapped reads, may result in falsely predicted alternative genotypes in multiple samples, thus representing false positive variant sites. In individual samples, false positive variants are manifested by falsely predicted alternative genotypes. Importantly, in WGA data, systematic errors may not be observed in all samples due to allelic dropout. To study the ability of Conbase and Monovar to detect false variants, we simulated a dataset with samples affected by errors. Because Conbase requires that at least two samples display reads supporting concordant alleles, we added two mock samples with heterozygous genotypes in all sites. The true genotype in all sites in all samples affected by increasing error rate, was defined as the reference genotype, e.g. no true mutation is present in the dataset. As such, we could determine the fraction of false positive genotypes by the number of alternative genotypes per sample. In samples affected by errors, no sites with falsely predicted alternative genotypes were present in the output from Conbase (Figure 2e, Supplementary Figure 2). In Monovar output, we observed sites affected by errors, to be incorrectly predicted as the alternative genotype. The fraction of false positive genotype calls per sample correlated with error rate (Figure 2e). The sites affected by errors exhibit support for alleles that cannot simultaneously exist, since a single allele defined by the gSNV cannot contain reads representing both mutant and reference genotypes at the putative sSNV site (assuming a diploid genome). Such observations are classified as conflicting genotypes by Conbase and can be distinguished from heterozygous genotypes with reads originating from concordant haplotypes (Figure 2d, Supplemenary Figure 2). Sites in which any sample in the dataset display support for discordant haplotypes are by default filtered out as low confidence variant sites by Conbase, instead of producing false positive variant calls (Methods: Algorithm description). All sites where errors were simulated in the dataset, where assigned ‘PASS’ in the variant site quality field (FILTER) in the Monovar variant calling format (vcf) output (Supplementary Figure 2).

### Performance evaluation on real world data obtained from *in vitro* expanded human fibroblasts

To evaluate the performance on real data, we first made time-lapse recordings of primary human fibroblasts as they divided on polymer slides allowing us to subsequently identify and isolate single fibroblasts with known phylogenetic relationships by laser capture microscopy. We isolated 11 cells derived from clone 1, three cells derived from clone 2 and two unrelated cells. All these single cells were whole genome amplified by multiple annealing and looping based amplification cycles (MALBAC)^12^. The single cell libraries were sequenced to obtain an average of 385 million reads per single cell, corresponding to approximately 12x coverage of the human genome on unamplified genomic DNA (Supplementary Table 1). We first estimated amplification efficiency relative to allelic dropout and locus dropout, by analyzing the fraction genomic bases covered by reads and the fraction of gSNV sites covered by reads originating from the maternal allele, the paternal allele or from both alleles (Supplementary Figure 3). On average, 26% of genomic bases were covered by at least one read, with a 70% allelic dropout rate at the covered gSNV sites (Supplementary figure 3). The low coverage and high allelic dropout observed in the MALBAC libraries is likely to reflect the fact that the cells were harvested by laser capture microscopy and thus part of the genomic material may be lost in the isolation process.

We next performed variant calling on bulk genomic DNA and single fibroblast libraries using FreeBayes^8^ and computed the fraction of sites in which an alternative genotype was observed in single cell libraries but not in the bulk sample (Supplementary Figure 3). These sites include a combination of true sSNVs and false positive variant calls. The unexpectedly high number of variants uniquely called in single cells (551971-1220408 unique variant calls per single cells), is suggestive of the presence of a large number of false positive variant calls. This is expected as MALBAC is reported to have a relatively high error rate due to the lack of proofreading by the *Bst* polymerase in the initial amplification steps, coupled with exponential amplification in the final steps of the protocol^9^. Moreover, variant callers designed for bulk data, including FreeBayes, do not account for the unique properties of WGA amplified single cell data, and may result in inaccurate SNV calling^4,5^.

We next performed variant calling with Monovar and Conbase, which are designed to account for the errors and biases in WGA amplified single cell data. To estimate the specificity of these methods to predict genotypes in real data from single cells, we computed the fraction of sites in which the distribution of genotypes was biologically plausible and implausible in our clonal populations of fibroblasts. True sSNVs are expected to be shared by closely related clonal cells, and not distributed between cells of different clones. Under the assumption that the probability of two mutations occurring independently in the same site twice is extremely low^10^, we defined implausible genotype distributions as sites where a variant call was observed in both clones and at least one cell displayed the reference genotype. Variants that are restricted to a single clonal population represent a biologically plausible genotype distribution. On raw Monovar output, we applied the recommended filtering^5^, including removal of sites overlapping with raw variant calling output of a bulk sample (obtained by FreeBayes), as well as sites present within 10 bases of another site. Parsing putative sSNVs from raw Monovar output yielded an unexpectedly high number of sites (Figure 3). To obtain only high confidence genotypes from Monovar output, we applied filters on the genotype quality (GQ) field on the variant calling output. GQ is a score calculated per sample by Monovar, reflecting the probability that the genotype prediction is correct. Low GQ scores indicate decreased confidence in the predicted genotype and high values indicate increased confidence in the predicted genotype. The genotypes in sites for individual samples that did not pass GQ score cutoffs were defined as unknown. Applying GQ filters resulted in a substantial decrease in the total number of called sites passing filters (Figure 3b, Supplementary Table 2). The majority of sites in Monovar output, displayed implausible genotype distributions, regardless of filters for the GQ and read depth (DP) fields, and for requirements that variants be observed in increasing number of samples (Figure 3). Following variant calling with Conbase, no further quality filtering is required and called genotypes can be used directly in downstream analysis. The fraction of implausible genotype distributions was small in Conbase output on MALBAC amplified fibroblasts, if a variant was required to be observed in >3 cells (Figure 3).

**Figure 3.**
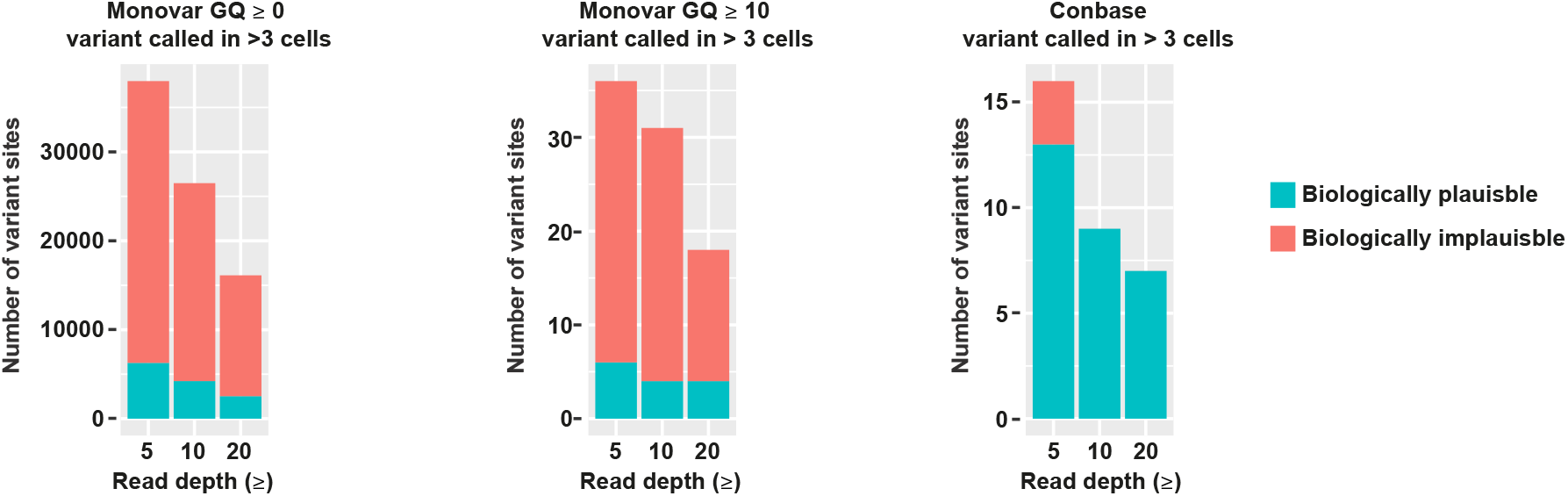
Biologically plausible and implausible distributions of reference and alternative genotypes called by Monovar and Conbase in clonal populations of fibroblasts. Biologically plausible genotype distributions were defined as sites where the variant call is exclusively observed within cells belonging to the same clone. Biologically implausible genotype distributions were defined as sites where the variant call is observed within both clones and at least one cell displayed the reference genotype.

### Performance evaluation on real world data obtained from *in vivo* expanded human CD8^+^ T cells

We next generated a dataset using multiple displacement amplification (MDA), a WGA method using the proofreading polymerase Phi29, which is associated with lower error rate and increased amplification efficiency^6^.

To evaluate Conbase on data generated from cells harvested directly from healthy human subjects, we examined CD8^+^ T cell clones, that had been expanded *in vivo* after yellow fever virus vaccination (YFV-17D). Clonally related cells were defined by sequencing of genomically rearranged T cell receptor (TCR) genes. Vaccination triggers the activation of individual naive T cells in lymphoid organs, leading to their expansion to large numbers of effector cells in the lymphoid tissues, which subsequently enter the circulation where they are detectable for at least several months after vaccination^12^. Expanded virus-specific CD8^+^ T cells can be identified and sorted from peripheral blood by labeling cells with fluorescently-labeled dextramers which are streptavidin linked HLA Class I complexes bound to a single viral epitope (Supplementary Figure 4)^11^. For this study we selected CD8^+^ T cells which had responded to a previously identified HLA-B7-restricited viral epitope for YFV (HLA-B7:RPIDDRFGL), which we observed to exhibit a reduced diversity of responding cells relative the dominant HLA-A2 epitope^12,13^. Single CD8^+^ T cells sorted by fluorescence-activated cell sorting (FACS) from longitudinal peripheral blood samples were amplified by MDA and subsequently screened by PCR against either the TCR a and/or β chains to identify clonally related T cells sharing the same TCR. Libraries from two clones, clone A (7 single cells) and clone B (9 single cells), as well as two unrelated cells (UR) were subjected to whole genome sequencing (Supplementary Table 1). As compared to MALBAC data, the percentage of bases covered by reads was higher in MDA data and the error rate was lower (Supplementary Figure 5)

Following variant calling with Conbase, no further quality filtering is required, although cutoffs for read depth were evaluated. As expected, decreased read depth cutoffs resulted in increased number of variant sites passing filters (Supplementary Table 2). Again, we compared our results with Monovar as a reference for the accuracy of Conbase. We attempted a range of cutoffs for DP and GQ, as well as requiring for a variant to be observed in increasing number of samples, in order to obtain putative sSNVs from Monovar output. Decreased DP cutoffs resulted in increased number of variant sites (Supplementary Table 2). In agreement with results from fibroblast data, applying no filters for GQ resulted in an unexpectedly high number of variant sites, while applying GQ cutoffs on Monovar output resulted in a substantial decrease of sites passing filters (Supplementary table 2).

To investigate if variants called by Conbase and Monovar represent true sSNVs, we performed unsupervised hierarchical clustering using shared genotype calls in sSNV sites to define distances between cells (Figure 4). Pairwise distances between cells were only based on sites where both cells had a genotype call. For sites where cells shared genotypes, the distance was decreased with −1, and increased with +1 if the genotypes differed. The distance matrix was then clustered using hclust with the distance method ward.D2. Hierarchical clustering based on genotypes called by Conbase demonstrated unambiguous identification of each T cell clonal population, regardless of read depth cutoffs (Figure 4a, Supplementary Figure 6). For Monovar output we attempted a range of combinations of filters to parse putative sSNVs from the vcf output. However, no combination of cutoffs for DP, GQ or when requiring for a variant to be present in increasing number of samples, revealed the two clonal populations in the dataset (Figure 4b, Supplementary Figure 7).

**Figure 4.**
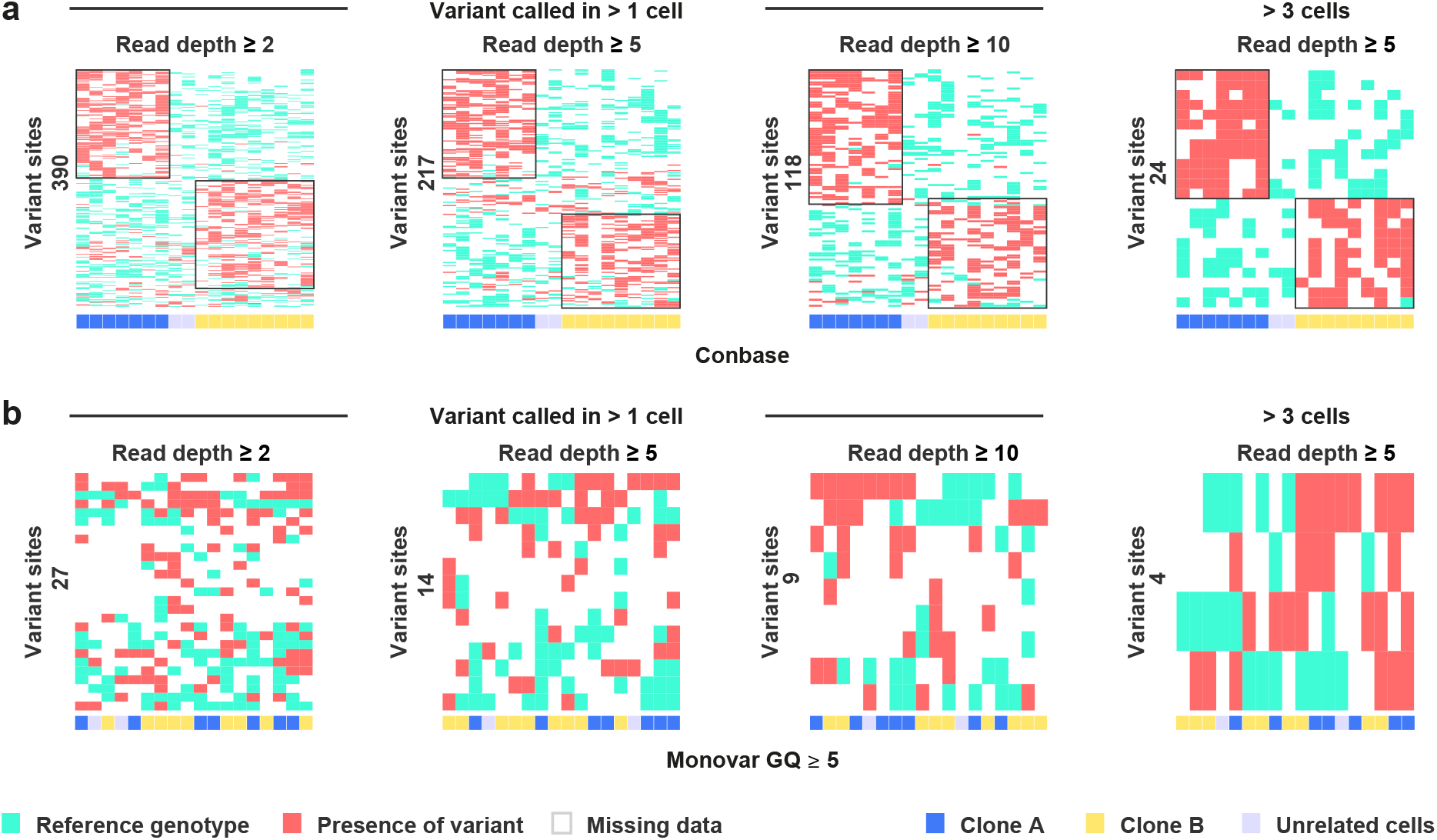
(a-b) Hierarchical clustering using genotypes called by Conbase and Monovar (GQ >= 5) to define distances between single T cells.

To estimate the fraction of false positive variant calls in the T cell dataset, we computed the fraction of biologically plausible (the variant is observed within a single clone) and implausible distribution of variant calls (the variant is observed in both clones and at least one cell is unmutated) in our clonal populations. The fraction of sites displaying biologically implausible genotype distributions in the T cell clones was small in Conbase output (Figure 5a). By default, Conbase filters sites in which at least one sample display conflicting genotypes, with support for both a mutated and an unmutated genotype. When allowing samples with conflicting genotypes in the final output from Conbase, we observed that 90% of sites in which at least one sample displayed conflicting genotypes also displayed a biologically implausible distribution of genotypes (data not shown). The vast majority of variant sites predicted by Monovar displayed implausible genotype distributions, independently of a range of combinations of cutoffs for DP, GQ and when requiring for a variant to be present in increasing number of samples (Figure 5b-d).

**Figure 5.**
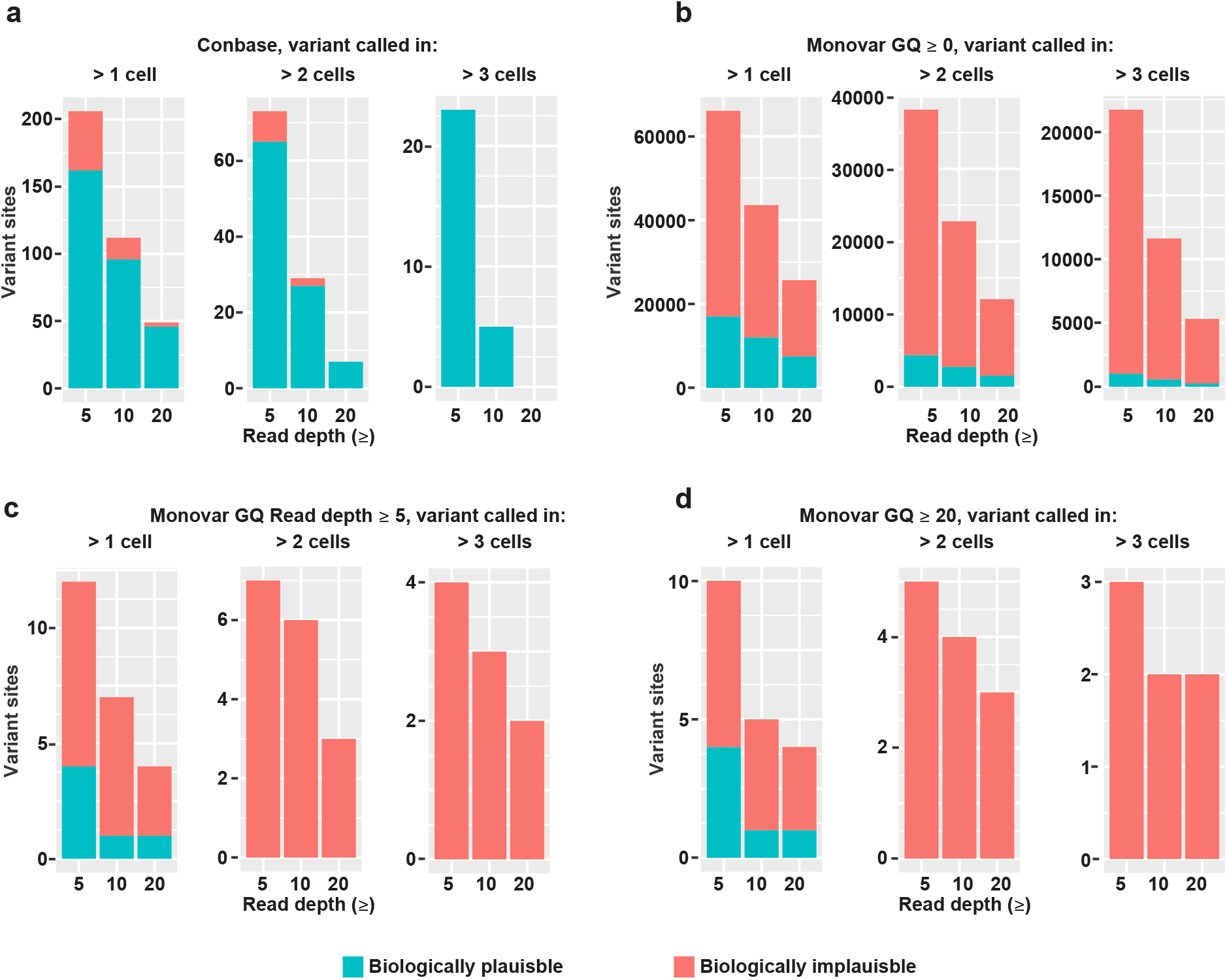
(a-d) Biologically plausible and implausible distributions of reference and alternative genotypes called by Conbase and Monovar in clonal populations of T cells. Biologically plausible genotype distributions were defined as sites where the variant call is exclusively observed within cells belonging to the same clone. Biologically implausible genotype distributions were defined as sites where the variant call is observed within both clones and at least one cell displayed the reference genotype.

### Validation of sSNVs Identified by Conbase

We validated a selection of the sSNVs called in T cells by Conbase, through PCR screening of additional MDA libraries generated from single CD8^+^ T cells isolated in parallel and determined to be clonally related by TCR sequence to the cells subjected to high-coverage whole genome sequencing. As a control, we included single CD8^+^ T cells identified as a third clonal population (Clone C, Supplemental Table 1) that was not included in the original Conbase screen. We designed PCR primers flanking regions containing both gSNVs and sSNVs in order to determine whether allelic dropout or amplification bias led to loss of informative alleles in our PCR products. Gel purified PCR amplicons from each single cell were subsequently subjected to Sanger sequencing to establish the presence or absence of the informative allele (gSNV) and candidate variant (sSNV). Here we could confirm the presence of sSNVs in cells belonging to the same clones, but never in unrelated cells, providing definitive evidence that the sSNVs called by Conbase represent true somatic mutations in each clonal lineage *in vivo* (Figure 6). In the Sanger results, we observed two sSNV sites (3:106210015 and 8:55449214), displaying heterogeneity within the clones, with some cells harboring the reference genotype and some cells harboring the alternative genotype. This could indicate that these two mutations appeared later during clonal expansion.

**Figure 6.**
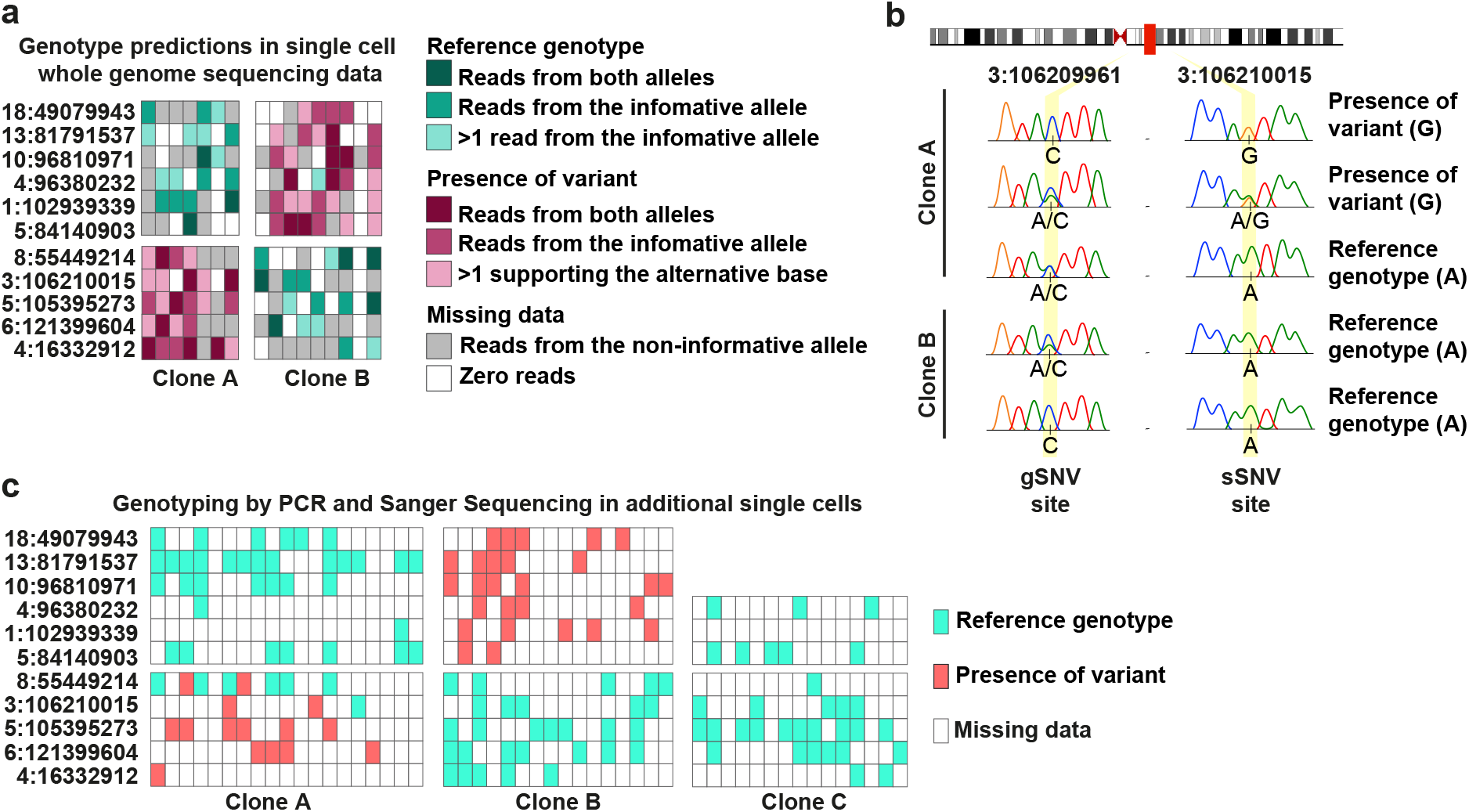
(a) Genotype predictions by Conbase in eleven sSNV sites. Primers spanning sSNVs and gSNVs were designed against these sites for PCR screening and Sanger sequencing. (b-c) Sanger Sequencing results for the selected sSNV sites in amplified DNA from single CD8^+^ T cells identified by TCR sequencing as belonging to clones A, B, or an unrelated clone C.

## Discussion

Conbase is to our knowledge the first available software capable of leveraging data from multiple samples in a dataset and utilizing read phasing to call and determine the presence or absence of somatic mutations in single cell WGA DNA libraries. Variant calling and genotyping is based on analysis of observed haplotype concordance, enabling exclusion of false positive variants resulting from WGA errors as well as systematic errors resulting from alignment artifacts. False negative genotype calls are eliminated through analysis of locus specific allelic dropout, determined individually per sample. As such, Conbase analysis is not dependent on global estimates of allelic dropout and genotypes can be predicted in samples exhibiting high rates of allelic dropout.

We have compared Conbase to Monovar on real and simulated data, as Monovar is the most widely used variant caller for single cell DNA sequencing data exhibiting allelic dropout and amplification bias^4^. On simulated data, Conbase achieves increased specificity over Monovar to correctly predict genotypes in the presence of allelic dropout and errors. The simulated data used in these experiments represent the errors and biases expected in WGA data from single cells, namely allelic dropout, amplification bias and errors deriving from amplification steps as well as alignment artifacts.

In real data, genotypes predicted by Conbase enable separation of clonally related populations of single cells from human donors, indicating that the called sSNVs represent true somatic mutations. Separation of the clonal populations was not achieved by genotypes predicted by Monovar, regardless of a range of combinations of cutoffs for DP, GQ and requiring for a variant to be observed in increasing number of samples. The failure to identify the clonal populations by hierarchical clustering may be attributed to the large fraction of false positive and false negative genotype calls observed in Monovar output as was seen in simulated data affected by allelic dropout and errors.

We show that the majority of variant sites called by Conbase in real data display biologically plausible distributions of genotypes in our clonal populations of cells, further indicating that variants called by Conbase represent true somatic mutations. In contrast, the majority of variant sites called by Monovar, display biologically implausible genotype distributions, indicating the presence of false positive variant calls. The small fraction of implausible genotype distribution in Conbase output as compared to Monovar output may be attributed to the ability of Conbase to detect and filter sites in which samples display conflicting genotypes, as investigated in simulated data affected by errors. In our real datasets, we observed that filtering sites displaying conflicting genotypes also removes presumably false positive variant calls, here represented by sites displaying biologically implausible distributions of genotypes. False positive variant calls are still present in Conbase output, albeit to a lower degree as compared to Monovar. These may be represented by false positive variant sites, in which no sample display conflicting genotypes, and can thus not be distinguished from true sSNVs.

Following variant calling with Conbase, we were able to confirm that identified sSNVs represent true somatic mutations by PCR screening of additional single cells sorted from the same donor and identified as being clonally related to the two clonal populations used for whole genome sequencing. We did not detect these variants in any cells isolated in parallel from a third unrelated clone. Indeed, we believe that this approach will provide a useful platform for expanding the analysis to hundreds or thousands of cells using targeted screening after identification of high confidence mutations in single cell whole genome sequencing data by Conbase.

## Conclusion

We report the development of a software that enables identification of somatic mutations at low read depth in single cell whole genome sequencing data exhibiting high rates of allelic dropout.

## Methods

### Algorithm description

Conbase requires whole genome sequencing data from WGA amplified single cells and an unamplified bulk sample to predict sSNV sites and genotypes. Conbase takes three inputs: single cell and bulk bam files, a human reference genome in fasta format and gSNV coordinates to be used for phasing. gSNV coordinates and gSNV base observations are obtained from vcf output previously generated from variant calling in a bulk sample by FreeBayes (or another variant caller). The analysis is split up in two subprograms: stats and analyze. Stats outputs a json-file with unfiltered variant calls, which is used as input to analyze. The output of analyze are phased filtered variant calls. Conbase variant calling is based on assumptions associated with expected observations in true sSNVs, including concordance of base observations in independent positions in reads and read pairs within samples and across the dataset (Supplementary Figure 8). The assumptions are coded as adjustable parameters with values reflecting how much read observations may deviate from the expected attributes of a true variant site. The parameter values generating the data presented in the current report are specified in parentheses in the algorithm description below.

#### stats

The initial step consists of identifying sites supporting an alternative (potentially mutated) base as compared to the bulk sample. This is done by only considering bases present in the same read or read pair as at least one gSNV using a BAM reader (pysam). The genomic windows in which the analysis will take place are determined by defining the longest distance upstream and downstream of each gSNV, covered by read pairs within the dataset. Because of sequencing errors, amplification errors or alignment artifacts, more than one alternative base may be observed in the same position, despite a diploid genome. In order to determine the most probable alternative base in a site, we utilize the accumulative information given by all qualified samples in the following way:

Let *R* be the reference base in the bulk and let *b* ∈ *B* = {*A, C, G, T*}\{*R*}. Let 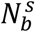 be the read depth of *b* in sample *s* at a given position and let and 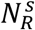 be the read depth *R*.

Let 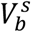 define the voting result for a base *b* by a sample *s*

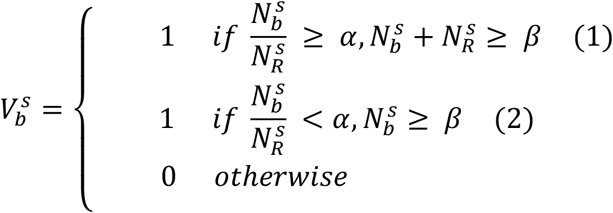

where *α* (0.2) and *β* (10x) are parameters for variant allele fraction and total read depth, determining which samples are qualified to vote. In the first case (1), reads from two alleles are observed, displaying sufficient variant allele fraction and total read depth. In the second case (2), the variant allele fraction is not sufficient, however the sample displays sufficient read coverage supporting the alternative base, which represents another indication for a potentially true variant since amplification bias may result in dropout of the reference base.

Let *V_b_* be the number of samples that voted for an alternative base *b*. If *V_b_* > 0 we may define the most probable alternative base *A* as

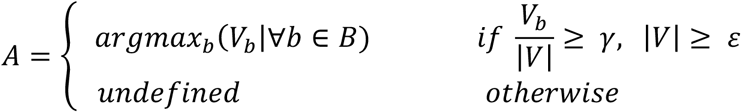

where *γ* (0.9) and *ε* (2) define the fraction of samples required to vote for the same base and the number of samples required to vote at a given position, respectively. |*V*| is the number of samples that were qualified to vote.

#### analyze

The previous processing identified potential sSNV sites in near distance of gSNVs where multiple cells shared strong support for the same alternative base *A*. If we consider the alternative base *A* in the sSNV site in relation to the base observed in a gSNV site in the same read or read pair, we denote it as the following relation:

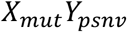

where the sSNV site *X_mut_* either displays the same base observed in the reference genome (and the bulk sample) or the alternative base, and *Y* either displays the same base as the reference genome or the alternative base in the gSNV site (obtained from vcf output from variant calling in the bulk sample). For simplicity, we will further denote these relationships as *RR,RA,AR,AA* where the first letter refers to the observation in the sSNV site and the second letter refers to the observation in the gSNV site (in both cases, we let *R* and *A* denote the reference and the alternative base, respectively). From here on we will denote a given pair as a *tuple*. Furthermore, a tuple pair, *tp*, denotes the possible combination of tuples on a chromosome pair.

Considering both the maternal and paternal pair, there are only two plausible tuple pairs that could equal a true heterozygous genotype. Likewise, there is only one single type of tuple pair that constitutes a true homozygous genotype. Hence, only three tuples are relevant, namely

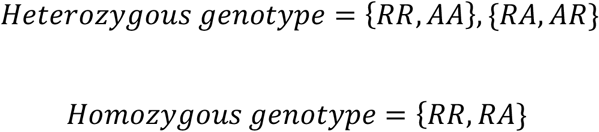

Given that we expect a clonal mutation to be present in a subpopulation of the analyzed cells, we can collectively determine the expected coupling representing a true heterozygous genotype, either {*RR, AA*} or {*RA, AA*}, by analyzing tuple pairs in all the samples that qualifies for voting.

All samples that display reads covering the tuple (i.e. the reads contain the specific gSNV in mind and the sSNV) with a total read depth > *φ* (see individual figures for the examined read depth cutoffs, Figure 3-5, Supplementary Figure 6) are allowed to vote if they fulfill the following criteria: Let *tp_max_, tp_min_* be the tuple pairs with the highest and lowest depth out of the two possible tuple pairs for heterozygous positions. A sample is allowed to vote for *tp_max_* if i) the ratio between the read depth of *tp_max_* and *tp_min_* < *ρ* (0.01) and ii) either the ratio for the tuples in *tp_max_* > *λ* (0.1) or if the tuple covering the variant {*AR or AA*} is the maximum out of all tuples for that sample. Out of the entitled samples that voted for their local *tp*_s_, the most probable tuple pair across all samples *tp*^*^ is finalized if at least *κ* (see individual figures for the examined number of samples required to harbor the variant, Figure 4–5, Supplementary Figure 6) samples voted and if *tp*^*^ held a majority of at least *ω* % (90%) of the votes. If the voting samples do not manage to conclude the final tuple pair for a sSNV in connection to a specific gSNV, that will dictate across all samples, the allelic origin of the alternative base is simply unknown.

By knowing the designated *tp*^*^ we also know which of the two alleles we expect to observe the variant on, either {*AA*} or {*AR*}. Consider the following: while we may jump to the conclusion of having found a variant as soon as we observe e.g. tuple *AA* in a sample, if the *tp*^*^ actually consists of tuples {*AR, RA*} we actually know for certain that this cannot be a true variant as such observation would be contradictory. Moreover, given this *tp*^*^, if a sample only display reads supporting the tuple *RA* we obviously cannot determine the genotype since RA is expected to be observed in both heterozygous and homozygous genotypes.

Let *N_x_* be the read depth of the tuple *x* and *N_tot_* the total read depth for all tuples. Consider the case when *tp*^*^ = {*RR,AA*} and let

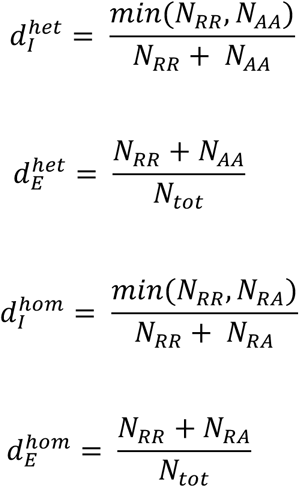

where 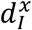 is the (internal) ratio for the tuples and 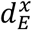 is the (external) ratio for a given tuple pair in relation to all of the tuples for a site from a homozygous and heterozygous perspective. These ratios will be used to determine the confidence in the allelic origin of the sSNV for a given site. The case when *tp*^*^ = {*RA,AR*} is treated analogously.

If *tp*^*^ is {*RR,AA*}, reads displaying *A* in the gSNV site is the informative allele, and reads originating from the informative allele are required for genotyping. If *tp*^*^ is {*RA,AR*}, reads displaying *R* in the gSNV site is the informative allele.

Genotypes are classified as being supported by reads originating from both alleles, or supported by reads originating from only the informative allele. This enables genotyping despite 100% allelic dropout, if reads originating from the informative allele are observed.

A genotype determination 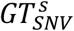 for any of these two classes of genotypes (either supported by reads originating from two alleles, or only supported by reads originating from the informative allele) for a given position in relation to a specific gSNV for a sample *s* will only be possible if there is a given *tp*^*^ and total read depth of all tuples > *φ* (see individual plots for the examined read depth cutoffs, Figure 3–5, Supplementary Figure 6). If these initial conditions are met, 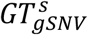 is computed as follows:

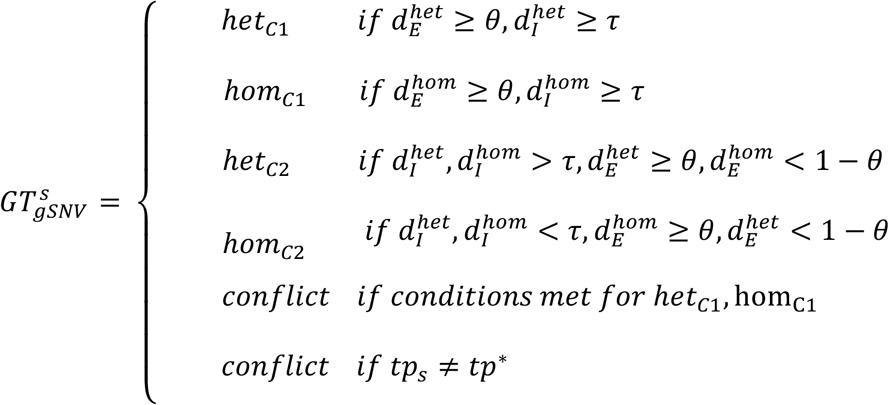

where *τ* (0.1) is the internal ratio parameter and *θ* (0.9) is the external ratio parameter.

The reads covering the sSNV may cover multiple gSNVs, which is why a collectively decided final genotype can be defined by letting all qualified 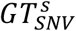 vote for the final genotype. *GT-1ax* is therefore the genotype with the most votes. Let *N_max_* be the number of gSNVs that voted for 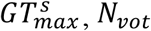 is the number of gSNVs that were qualified to vote (*tp*^*^ is defined for this gSNV). Let *N_tot_* be the number of gSNVs that were present in the same read or read pairs as the sSNV and let *GT^S^* be defined as

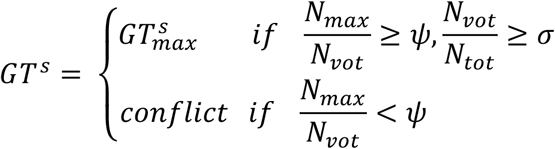

where *ψ* (0.9) is the fraction of gSNVs voting for the same genotype, *σ* (0.9) is the fraction of gSNVs qualified to vote for a genotype.

If *N_vot_* = 0, any available read information is used to infer genotypes for samples displaying insufficient read depth. Support for an unmutated genotype is defined as when either the *tp*^*^ = {*AA,RR*} and the sample displays a read depth supporting the tuple *RA* > *χ* (1x) or when the *tp*^*^ = {*AR, RA*} and the sample displays a read depth supporting the tuple *RR* > *χ*, resulting in *GT^s^* = *hom*_*C*3_. If there is no support for an unmutated genotype and the sample displays a read depth supporting the alternative base > *ξ*(1x), *GT^s^* = *het*_*C*3_. Samples displaying support for both a mutated and an unmutated genotype are considered *GT^s^ = conflict*.

In the current report zero samples displaying conflicting genotypes were allowed per site. A common artefact in variant calling output, is regions with clusters of false positive mutations, correlating with areas in the genome regions with poor mappability. In the current report, a maximum of 1 mutation per kb was allowed.

### Sequence alignment and data processing

Following whole genome sequencing and demultiplexing, the reads were trimmed from Illumina adapters using Cutadapt^14^. WGA adapters were trimmed from MALBAC amplified samples using Cutadapt^14^. Read pairs were aligned to the human genome (human g1k v37 with decoy) using Burrows-Wheeler aligner (BWA-MEM)^15^. Processing of the mapped reads and sequencing data quality evaluation was performed using Picard Tools (http://picard.sourceforge.net)and FastQC (http://www.bioinformatics.babraham.ac.uk/projects/fastqc/). Read processing included removal PCR duplicates, optical duplicates and reads with a mapping quality below 2, including multimappers. Indel realignment was performed with GATKs IndelRealigner^16^. Variant calling was performed using FreeBayes with default settings^8^.

### gSNV filtering

Following variant calling in bulk samples using FreeBayes, variants were filtered by vcffilter (https://github.com/vcflib/vcflib) to identify gSNVs (Supplementary Figure 9). False gSNVs will result in false variant calls in the downstream analysis with Conbase. As such, we applied stringent filters on the vcf output from FreeBayes. With an average read depth of ≈40x in our bulk samples, we estimated a conservative maximum depth threshold of 55x at any position, as

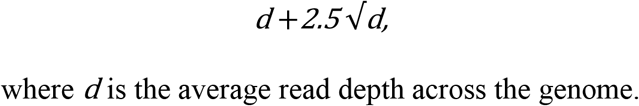

where *d* is the average read depth across the genome.

We filtered variants on autosomes with a 15-55-fold read depth, quality score above 10 (QUAL), reads originated from both strands (SAF & SAR), at least two reads were balanced on each side of the site (RPR & RPL) and the alternative allele observation count was required to range between 20%-80% (AO). The variants were further filtered to remove gSNVs present in suspected erroneous regions in the reference genome. This was done by running a separate script included in the Conbase package, which screens the bulk bam file around each gSNV, and excludes gSNVs present within a 1kb window containing >10 ‘heterozygous’ positions. A ‘heterozygous’ position was defined as a position where >10% of the reads supported a non-reference base. In the fibroblast donor, 1634933 gSNVs passed filters. In the T cell donor, 1789830 gSNVs passed filters. ~70% of gSNVs are present within 1200 bases of another gSNV (Supplementary Figure 9). ~50% of genomic bases are present within 600 bases of a gSNV and can thus be phased if the insert size of the sequencing library is 600 bp (Supplementary Figure 9).

### Generation of Simulated Data

To generate data with known genotypes we selected gSNV sites in a bulk sample from the T cell donor (filtered as above) present 15 bases apart, where the alternative base in the downstream gSNV site was present on the same allele as the alternative base in the upstream gSNV site. We next filtered these loci to obtain sites were the Fibroblast donor (bulk sample) was heterozygous in the downstream gSNV site, and homozygous for the reference base in the upstream site. A total of 40 loci were passed these requirements, representing one gSNV site and one neighboring simulated variant site. Read data covering these loci were extracted by samtools view. Data from the T cell donor represent a sample with reads from both alleles mutated in all sites (sample_mut). Data from the Fibroblast donor represent a sample with reads from both alleles unmutated in all sites (sample_unmut). To simulate allelic dropout data that would be representative of the read depth distributions observed in WGA data, we extracted reads from single cells with coverage over the simulated loci. We constructed one file containing reads only displaying the reference base in the (downstream) gSNV site (sample_ADO_ref), and one file only containing reads displaying the alternative base in the (downstream) gSNV site (sample_ADO_alt). Samples affected by allelic dropout were simulated by extracting and merging reads from sample_mut, sample_ADO_ref and sample_ADO_alt (using samtools). At 50% allelic dropout, 50% randomly selected sites were covered by reads extracted from sample_mut. Of the remaining sites, 50% randomly selected sites were covered by reads extracted from sample_ADO_ref and the remaining sites were covered by reads extracted from sample_ADO_alt. Samples affected by errors were simulated by merging reads from sample_unmut and sample ADO_alt in increasing number of sites.

### Clonal human fibroblast Isolation and Analysis

Single cells isolated from a primary human fibroblast cell line C5RO (normal) were expanded *in vitro* on a Leica frame slide. Clonally related cells (determined by time lapse movie recording) were isolated by LCM. 11 cells from clone1, three cells from clone2 and two unrelated cells were next subjected to WGA using MALBAC (Yikon Genomics). Samples were individually inspected using a bioanalyzer (Agilent) and library preparation was done using KAPA HTP Library Preparation kit Illumina Platform (KR0426,KAPABI0SYSTEMS) and whole genome sequencing. The cells belonging to clone 1 were sequenced to an average depth of 15x. The single cells belonging to clone 2 and unrelated cells were sequenced to an average depth of 10x. An unamplified bulk sample from the same primary cell line was sequenced to an average depth of 40x.

### T cell Sample Preparation and Cell Sorting

Study participants were recruited into an ongoing study to monitor immune responses to the yellow fever virus vaccine YFV-17D (approved by the Regional Ethical Review Board in Stockholm, Sweden: 2008/1881-31/4, 2013/216-32, and 2104/1890-32). A female subject was identified based on being positive for HLA-B7 and having a detectable T cell response to a minor peptide (RPIDDRFGL) presented by HLA-B7. Cryopreserved peripheral blood mononuclear cell (PBMC) samples taken at days 10, 30, and 148 post-vaccination were thawed at 37°C and quickly washed in FACS buffer (PBS with 2% BSA/2mM EDTA). Negative selection with magnetic beads was performed for each sample to purify CD8^+^ T cells (Miltenyi Human CD8 Negative Selection kit, 130-096-495). Purified CD8^+^ T cells were first incubated with an HLA-B7/RPIDDRFGL dextramer conjugated to Alexa fluor 647 (Immudex) for 15 minutes. Cells were subsequently incubated with a panel of antibodies to identify live CD3^+^CD8^+^ T cells (CD3–Alexa Fluor 700 (UCHT1, BD Biosciences), CD8-APC-Cy7 (SK1, BD Biosciences), CD4-PE-Cy5 (RPA-T4, eBioscience), CD14–Horizon V500 (MΦP9, BD Biosciences), CD19-Horizon V500 (HIB19, BD Biosciences), and Live/Dead Fixable Aqua Dead Cell staining kit (Invitrogen, L34957)). Live, Lineage negative CD3^+^CD8^+^Dextramer^+^ cells were sorted into 96 well PCR plates (Thermo Scientific, AB-0800) containing lysis buffer (200mM KOH, 40mM DTT, 5mM EDTA). Single cells were incubated on ice for 10 minutes in lysis buffer after which neutralization buffer (400mM HCL, 600mM Tris-HCL pH7.5) was added followed by an additional 10-minute incubation on ice. Lysed cells were subsequently stored at −80°C until amplification reactions were performed.

### WGA by MDA

Lysed single T cells were subjected to multiple displacement amplification (MDA) as previously described^17^. A mixture containing dNTPs (Invitrogen, 2mM), random hexamer primers with 3’ thiophosphate linkers (5’-dNdNdNdN*dN*dN-3’, IDT (50uM)), and repliPHI polymerase (40U) in phi29 reaction buffer (Epicentre) was added to each well to bring total volume to 20uL. Cells were incubated at 30°C for 10 hours followed by a 3-minute incubation at 65 °C to inactivate the phi29 polymerase. The resulting libraries were diluted in H2O to 50uL and concentrations of double stranded DNA were measured (Qubit, Broad Range dsDNA kit).

### Identification T Cell Receptors from Single Cell MDA Material

We adopted a previously published method^18^ so that we could screen large numbers of single cell libraries to identify clonally related T cells by TCR rearrangements. Approximately 100ng of amplified DNA was taken from each sample and touchdown PCR (Tm: 72°C -> 55°C) was performed using a panel of primers designed upstream of each variable region and downstream of the joining regions for the human TCR *a* or β chain locus (Supplementary Table 3). A dilution of each reaction was subsequently used to perform a second, nested-touchdown PCR with internal primers designed against each variable and joining region of the human TCR a or β chain locus. The internal primers contained handles which were used to index each well for the 96-well plate so that they could be pooled into a single reaction. Each plate was then prepared according to the Truseq (Illumina) protocol for sequencing on an Illumina Miseq (2×150bp reads). After demultiplexing of Illumina sample indexes, the reverse read (R2, 150bases) Fastq file was converted to Fasta format. Identical sequences were clustered using the FASTX-Toolkit (http://hannonlab.cshl.edu/fastx_toolkit/) FASTA Collapser. Then sequences were sorted by our 96-well indexes using the FASTX barcode splitter, and the first 44 bases were finally trimmed off using the FASTA trimmer to facilitate downstream sequence analysis. Because the internal primers targeting the joining regions were within 50bp of the CDR3 region of the TCR it was possible to identify clonal T cells based on shared CDR3 nucleotide sequences. All samples were individually analyzed using the IMGT database to identify the CDR3 sequence^19^.

### Selecting High Coverage Libraries for Illumina Sequencing

Clonal T cells were grouped and high quality libraries were identified using a panel of chromosome specific PCR primers as described previously^9^. High quality T cell libraries were considered to be samples with detection at the majority of loci and were subsequently processed using a PCR-free TruSeq library preparation kit (Illumina) and sequenced with a HiSeq X using a theoretical coverage of 30x per sample (SciLifeLab, Karolinska Institute).

### Screening Related Clonal T Cells by Sanger Sequencing

Single cell libraries that were included in the original screening which matched clones A or clone B were identified to be used for verifying selected mutants (summarized in Supplementary Table 4). An additional clone (Clone C (TCRa: CAAHSPYSGNTPLVF, TCRβ: CASSSGTAYNEQFF) was used as a control to determine whether mutations could be found as artifacts in unrelated T cells. Primers were designed to span both the gSNV and the putative variants and samples were subjected to 35 cycles of PCR (Tm: 67°C) (PCRBIO HiFi Polymerase, PCR Biosystems) yielding approximately 1000bp amplicons (Supplementary Table 4). Additionally, primers contained handles (similar to those used for TCR screening) so that secondary amplification cycles could be used to index samples if necessary. Amplified samples were analyzed by gel electrophoresis and bands were excised for DNA isolation (Nucleospin Gel Clean Up, Techtum). Gel-purified DNA samples were sent for Sanger sequencing (KI Gene Facility, CMM, Karolinska Institute) using primers specific for the universal handle incorporated onto each Forward primer (Supplementary Table 4). Sanger sequencing results were analyzed visually using the software package 4peaks and are summarized in Supplementary Table 4.

### Comparisons of single cell variant calling algorithms for performance evaluation

Monovar^5^ was run on single T-cell amplified with MDA with default settings including consensus filtering. From the raw Monovar output, sites present within 10bp of another site were removed. Sites overlapping with raw variants called in an unamplified bulk sample by FreeBayes, were removed. Potential sSNVs were filtered on autosomes by requiring that at least two samples shared a variant (0/1 or 1/1) while at least one sample displayed the reference genotype (0/0). Samples in the same site which did not pass these cut offs were assigned an unknown genotype. We attempted to call variants using SCcaller^6^. The rate of amplification bias observed in the T cell dataset and the Fibroblast dataset resulted in eta-values that were too low to enable distinction between true mutations and artefacts (personal communication with Xiao Dong^5^). Thus, we did not move forward with variant calling using the SCcaller.

### Hierarchical clustering

Variants called by Conbase and Monovar were used to define distances between cells. Distances between cells were defined as unknown if no shared sites were detected. For shared sites, the distance was decreased with −1 for each site where cells have the same call (mutated or non-mutated) and increased with +1 for sites where the cells have different calls. The distance matrix was then clustered using standard hclust with the distance ‘ward.D2’. For Monovar matrices with more than 45K sites (no applied GQ filtering), 45K randomly selected sites were included in the clustering analysis.

### Conbase Output

The final output from Conbase includes a tsv file with phased variant calls and a complementary interactive html file where genotype predictions are color coded based on presence or absence of mutation, presence or absence of allelic dropout, read depth support as well as summarized statistics about concordance of base observations in phased reads in the predicted variant sites for each sample.

## Abbreviations

WGA: Whole Genome Amplification
gSNV: Germline Single Nucleotide Variant
sSNV: Somatic Single Nucleotide Variant
MALBAC: Multiple Annealing and Looping Based Amplification Cycles
MDA: Multiple Displacement Amplification
DP: Read depth field in Monovar variant calling format output
GQ: Genotype quality field in Monovar variant calling format output
FACS: Fluorescent Activated Cell Sorting

## Declarations

### Ethics approval and consent to participate

All human subjects involved in this study provided written, informed consent affirming their participation in this study. The ethical permit was approved by the Regional Ethical Review Board in Stockholm, Sweden: 2008/1881-31/4, 2013/216-32, and 2104/1890-32

### Consent for publication

All human subjects involved in this study provided written, informed consent acknowledging that any findings arising during this study would be submitted for publication and data would be deposited upon publication.

### Competing financial interests

The authors declare no competing financial interests.

### Funding

This work was supported by the Swedish Research Council, the Swedish Cancer Society, the Swedish Society for Strategic Research, Tobias Stiftelsen, AFA Försäkringar, the Strategic Research Programme in Stem Cells and Regenerative Medicine at Karolinska Institutet (StratRegen), Torsten Söderbergs Stiftelse and the Swedish Genomes Program (Science for Life Laboratory). The Swedish Genomes Program has been made available by support from the Knut and Alice Wallenberg Foundation. Å.B. and B.S. are financially supported by the Knut and Alice Wallenberg Foundation as part of the National Bioinformatics Infrastructure Sweden at SciLifeLab.

### Author Contributions

J.H. and J.E.M devised the project. J.H., M.K. and E.A.H. developed the software, M.K and E.A.H implemented and J.H analyzed data. J.E.M. conceived and planned the experiments with support from I.D., M.P. and J.M. Å.B., contributed to data analysis with support from E.B., P.S., B.S. J.H., M.K., E.A.H., and J.EM. wrote the manuscript with contributions from the remaining authors. J.E.M. and J.F. supervised the project.

## Acknowledgements

J.H. would like to acknowledge bioinformatics advice through the Swedish Bioinformatics Advisory Program.

